# Deep mutational scanning and massively parallel kinetics of plasminogen activator inhibitor-1 functional stability

**DOI:** 10.1101/2022.07.19.500671

**Authors:** Laura M. Haynes, Zachary M. Huttinger, Andrew Yee, Colin A. Kretz, David R. Siemieniak, Daniel A. Lawrence, David Ginsburg

## Abstract

Plasminogen activator inhibitor-1 (PAI-1), a member of the serine protease inhibitor (SERPIN) superfamily of proteins, is unique among SERPINs for exhibiting a spontaneous conformational change to a latent or inactive state. The functional half-life for this transition at physiologic temperature and pH is ~1-2 h. To better understand the molecular mechanisms underlying this transition, we now report on the analysis of a comprehensive PAI-1 variant library expressed on filamentous phage and selected for functional stability after 48 h at 37 °C. Of the 7,201 possible single amino acid substitutions in PAI-1, we identify 439 that increase the functional stability of PAI-1 beyond that of the wild-type protein and 1,549 that retain inhibitory activity toward PAI-1’s canonical target protease (urokinase-like plasminogen activator, uPA), while exhibiting functional stability less than or equal to that of wild-type PAI-1. Missense mutations that increase PAI-1 functional stability are concentrated in highly flexible regions within the PAI-1 structure. Finally, we developed a method for simultaneously measuring the functional half-lives of hundreds of PAI-1 variants in a multiplexed, massively parallel manner, quantifying the functional half-lives for 697 single missense variants of PAI-1 by this approach. Overall, these findings provide novel insight into the mechanisms underlying PAI-1’s latency transition and provide a database for interpreting human PAI-1 genetic variants.

## INTRODUCTION

<u>S</u>erine <u>p</u>rotease <u>in</u>hibitors, or SERPINs, comprise a protein superfamily with over 30 members in humans, the majority of which display specific inhibitory activity towards a subset of serine proteases (1,2). Despite the highly-specific interaction of most SERPINs with their target proteases, all SERPINs share a common structure that consists of a globular domain containing a central β-sheet (β-sheet A), with a reactive center loop (RCL) that extends above the core of the protein when the SERPIN is in its metastable active state. The RCL contains a sequence motif that mimics the primary recognition site for its unique set of target serine protease(s) (3). When the serine protease encounters its target sequence in the RCL, proteolysis between the P1 and P1’ positions is initiated by the formation of an acyl intermediate. Inhibition of the target serine protease results when the RCL inserts into β-sheet A prior to hydrolysis of the acyl intermediate (4), with the resulting conformational changes rendering both the serine protease and SERPIN no longer functional. If the acyl-intermediate is hydrolyzed prior to insertion of the RCL into β-sheet A, the serine protease will remain active, while the proteolyzed RCL will render the cleaved SERPIN non-inhibitory (1,5).

Plasminogen activator inhibitor-1 (PAI-1, *SERPINE1*) is a 379 amino acid member of the SERPIN superfamily composed of nine α-helices and three β-sheets. PAI-1’s canonical function is to inhibit the serine proteases urokinase-like and tissue-type plasminogen activator (uPA and tPA, respectively)—the primary proteolytic activators of the zymogen plasminogen to the serine protease plasmin. Plasmin is the key enzyme responsible for fibrinolysis, the proteolytic degradation of fibrin clots. Although the major function of PAI-1 appears to be in hemostasis through regulation of the fibrinolytic pathway, other functional roles have been proposed (6). *In vivo*, PAI-1 circulates in an 1:1 complex with its cofactor vitronectin (7,8), which also enhances PAI-1’s ability to inhibit other proteases in the coagulation pathway including thrombin and activated protein C (9). PAI-1 has also been demonstrated to inhibit other coagulation factors including coagulation factor IXa, coagulation factor XIIa, plasmin, and kallikrein (8,10), as well as non-coagulation proteases including furin and TMPRSS2—both of which have been implicated in the priming of viruses for cellular entry, including influenza A and SARS-CoV-2 (11–15).

Mice deficient in PAI-1 are viable without hemostatic abnormalities (16,17), and PAI-1 deficiency has been reported to be protective in animal models for a number of disease states (18). Complete PAI-1 deficiency in humans results in a mild to moderate bleeding diathesis (19,20), with heterozygous PAI-1 deficiency linked to increased longevity (21). Elevated levels of plasma PAI-1 have also been associated with a number of disorders including myocardial infarction, atherosclerosis, fibrosis, obesity, type 2 diabetes, metabolic syndrome, and cancer (6,22,23).

PAI-1 is unique among the SERPIN protein superfamily in undergoing a spontaneous transition to a non-inhibitory but more thermodynamically stable, latent conformation when, in the absence of a proteolytic event, an intramolecular rearrangement occurs in which the RCL spontaneously inserts into β-sheet A. The metastable PAI-1 active conformation exists in equilibrium with a prelatent state in which the RCL is hypothesized to be partially inserted into β-sheet A (24–29). Full insertion of the RCL is conformationally blocked by the Val^157^-loop between helix F and β-strand 3A; however, partial unfolding of helix F permits full insertion of the RCL into β-sheet A, while the return of helix F to the native state traps PAI-1 in its latent conformation (26,27).

The capacity of PAI-1 to undergo this latency transition is conserved across vertebrates (30–32), with human PAI-1 exhibiting a half-life (t_1/2_) in the active conformation of approximately 1-2 hours (29,33–35), at physiologic temperature and pH. The half-life is modestly prolonged in the presence of vitronectin (29,36), which stabilizes PAI-1 in its metastable active conformation. While the physiologic function of the PAI-1 latency transition is unknown, it may serve to clear plasma PAI-1 activity from circulation that is distal from a site of fibrinolytic activity (37), as well as a regulatory mechanism to limit the high PAI-1 levels that are associated with several disorders and linked to poorer outcomes, as noted above. PAI-1 variants engineered to have extended half-lives have also been suggested as potential therapeutics to promote wound healing (38–40). Similarly, cataloging the functional stability and half-lives of PAI-1 variants will provide a useful database given the interest in engineering novel SERPINs as potential therapeutics as it would provide a mechanism to limit or extend bioavailability (41,42). In a previous report, we used deep mutational scanning (DMS) to characterize the impact of PAI-1 missense variants on uPA inhibitory activity (43). This study identified missense mutations in PAI-1 that are both tolerated and not tolerated with respect to uPA inhibition. In this study, we expand on these results by performing a deep mutational scan on PAI-1 latency to examine how mutations in PAI-1 modulate its transition into a latent non-functional state (43).

To date, a limited number of PAI-1 variants have been engineered with enhanced functional stability and extended half-lives, using both unbiased and rational design approaches (38,44–47); however, a comprehensive analysis of both gain- and loss-of-function variants with respect to functional stability has not been reported. We also now describe a novel approach to simultaneously measure the half-lives of multiple PAI-1 variants in a massively parallel fashion. Together these data provide an empirical method to assess allosteric networks within PAI-1 that contribute to its latency transition. Our findings also complement the molecular dynamics simulations of Kihn *et al* (48).

## RESULTS

### Massively parallel determination of PAI-1 variant functional stability

The fraction of active phage displayed PAI-1 (phPAI-1) (43) was determined by immunoprecipitation of phPAI-1:uPA complexes with an α-uPA antibody at the indicated time points of incubation at 37°C over a 48 h time course (Fig. 1). Using this approach, the half-life (t_1/2_) of WT phPAI-1 was determined to be 2.1 h, consistent with published values for WT recombinant PAI-1 (rPAI-1) (29,33,34). In parallel experiments, immunoprecipitation by the N-terminal *myc*-tag of phPAI-1 further confirmed that the loss of phPAI-1 functional activity is due to its transition to latency and not degradation or proteolysis (Fig. 1). When the previously characterized phPAI-1 mutant library (43) was analyzed over the same 48 h time course, the apparent aggregate half-life of the library was determined to be 11.2 h (Fig. 1). This prolonged half-life is likely explained by the presence within the library of multiple phage carrying mutations that prolong the functional stability of PAI-1.

**Figure 1.**
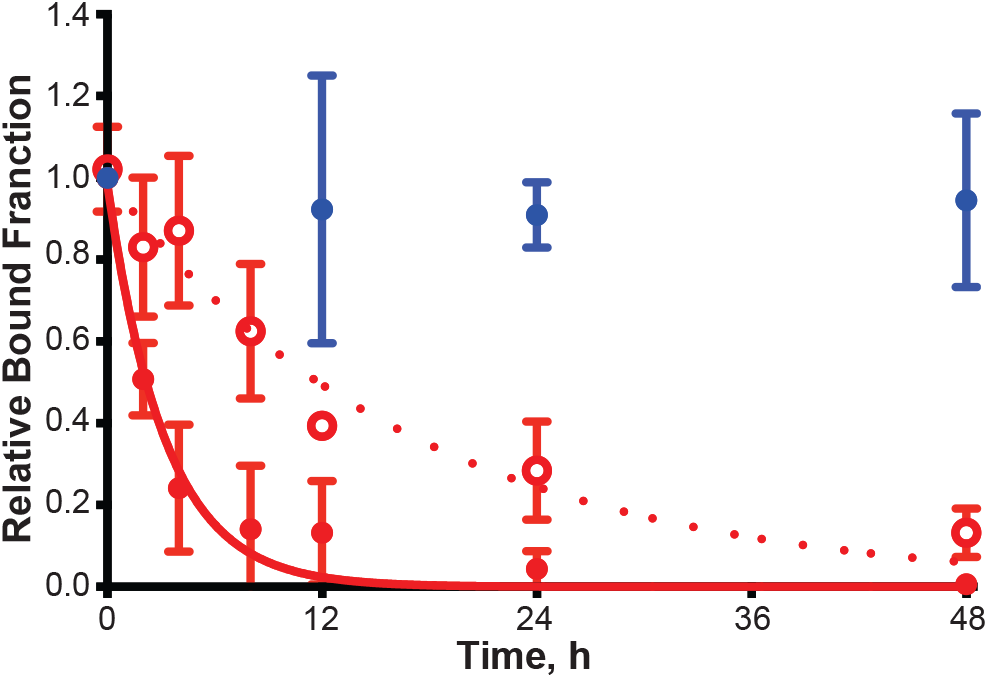
The phPAI-1 mutant library undergoes an extended latency transition compared to WT phPAI-1. The relative quantity of WT phPAI-1 (closed red circles) or mutant library phPAI-1 (open red circles) recovered by uPA complex formation and α-uPA immunoprecipitation at selected time points (0, 2, 4, 8, 12, 24, 48 h) at 37° C. The data were fit to a one-phase exponential decay function (Eq. 1) and the apparent half-lives of WT phPAI-1 and the mutant library were determined (Eq. 2). *Myc*-tag immunoprecipitation (closed blue circles) preformed at selected time points (0, 12, 24, 48 h) at 37° C shows no significant decrease in number of phage recovered after 48 h incubation. Mean phage ± SD relative to amount immunoprecipitated at time 0 is plotted. N=3 per group for all time points.

We previously reported the construction of a phPAI-1 mutant library containing 8.04 × 10^6^ independent clones with an average of 4.2 ± 1.8 missense mutations per fusion protein (43). To assess the relative functional stability of phPAI-1 variants that retained inhibitory activity at 48 h, the phPAI-1 mutant library was incubated at 37°C for 48 h before selection for complex formation with uPA (SI Data 1). Input and selected phage were analyzed by high throughput DNA sequencing (HTS). Variants that were non-functional at 0 h (i.e., depleted following selection for complex formation without heat-induced latency) were removed from subsequent analyses as these variants were assumed to either have no inhibitory activity or an extremely short functional half-life (43). The relative functional stability scores for missense mutations compared to the WT amino acid at each position are illustrated in Fig. 2. Of the 1,988 missense mutations that were characterized for functional stability relative to WT PAI-1, 439 amino acid substitutions generated PAI-1 variants with increased functional stability. Conversely, 1,549 amino acid substitutions resulted in PAI-1 variants that were functionally active yet less functionally stable than WT PAI-1. The missense mutations with the top 35 functional stability scores are listed in Table 1. Although PAI-1 mutations that increase or decrease the rate of uPA inhibition may contribute to relative functional stability scores, this effect should be negligible given the high uPA concentration compared to that of phPAI-1 in addition to the relatively long reaction time (30 min).

**Figure 2.**
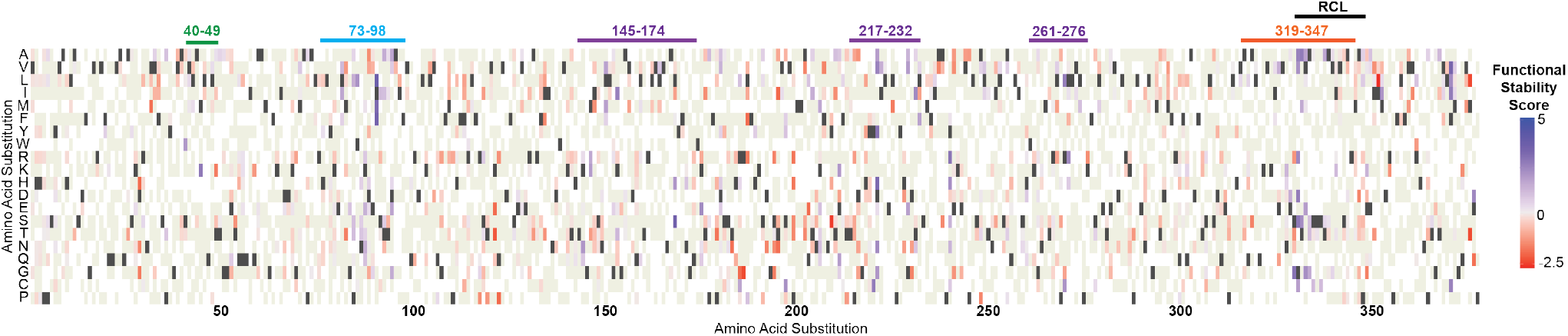
DMS identifies PAI-1 missense variants that exhibit inhibitory activity towards uPA with changes in functional stability. Heatmap indicating functional stability relative to WT PAI-1 for phPAI-1 variants selected for uPA inhibitory activity at 48 h. Amino acid position is indicated on the x-axis, while amino acid substitutions are indicated on the y-axis. Variants with increased functional stability relative to WT are indicated in blue, while those with decreased functional stability are highlighted in red. WT amino acid residues are shown in dark grey, while beige indicates missense mutations that were present in the mutational library but did not meet statistical thresholds (43). Regions enriched in variants exhibiting enhanced functional stability include the RCL (residues 331-350, *black*) as well as residues 40-49 (*green*), residues 73-98 (*cyan*), residues 145-174 (*purple*), residues 217-232 (*purple*). residues 261-276 (*purple*), and residues 319-347 (*orange*).

**Table 1.** 35 most functionally stable PAI-1 variants relative to WT

| Mutation | Functional Stability Score | p<sub>adj</sub> | Massively Parallel Half-life, h |
| --- | --- | --- | --- |
| I91L | 3.73 | $2.27 \times 10^{-188}$ | ND |
| L169S | 3.67 | $6.25 \times 10^{-29}$ | ND |
| I91F | 3.47 | $<10^{-294}$ | 7.7 |
| F372V | 3.39 | $<10^{-294}$ | 6.1 |
| D222H | 3.22 | $5.14 \times 10^{-136}$ | ND |
| I91M | 3.20 | $5.51 \times 10^{-165}$ | ND |
| S331G | 2.97 | $<10^{-294}$ | 6.8 |
| D222Y | 2.95 | $4.76 \times 10^{-293}$ | ND |
| Y241R | 2.94 | $2.66 \times 10^{-26}$ | ND |
| M45K | 2.82 | $<10^{-294}$ | 3.9 |
| G332S | 2.81 | $4.63 \times 10^{-242}$ | 6.2 |
| T333A | 2.74 | $<10^{-294}$ | 7.8 |
| Y221C | 2.72 | $5.47 \times 10^{-240}$ | ND |
| F372I | 2.72 | $<10^{-294}$ | 4.0 |
| G332D | 2.70 | $9.24 \times 10^{-225}$ | 7.0 |
| R271L | 2.67 | $3.20 \times 10^{-239}$ | 3.7 |
| K154I | 2.62 | $1.21 \times 10^{-266}$ | 3.3 |
| G332A | 2.62 | $2.48 \times 10^{-124}$ | ND |
| F372C | 2.57 | $8.19 \times 10^{-123}$ | ND |
| D222V | 2.56 | $1.54 \times 10^{-263}$ | ND |
| E378D | 2.45 | $7.76 \times 10^{-236}$ | 4.9 |
| G332R | 2.44 | $1.60 \times 10^{-55}$ | ND |
| T323I | 2.40 | $<10^{-294}$ | ND |
| L272R | 2.38 | $1.95 \times 10^{-193}$ | 3.4 |
| K176E | 2.38 | $1.26 \times 10^{-210}$ | 4.1 |
| D222N | 2.37 | $<10^{-294}$ | 2.0 |
| F372L | 2.34 | $<10^{-294}$ | 3.7 |
| T333S | 2.33 | $8.77 \times 10^{-65}$ | 8.4 |
| Y241H | 2.31 | $<10^{-294}$ | 2.0 |
| V334G | 2.28 | $1.37 \times 10^{-89}$ | ND |
| S198C | 2.26 | $4.44 \times 10^{-118}$ | ND |
| G332V | 2.18 | $1.80 \times 10^{-70}$ | 7.6 |
| W86L | 2.17 | $9.72 \times 10^{-274}$ | 13.1 |
| R271C | 2.16 | $4.50 \times 10^{-277}$ | 3.9 |
| L169H | 2.14 | $1.68 \times 10^{-88}$ | 3.2 |

p<sub>adj</sub> = adjusted p-value as defined by Love, *et al.* (61)
ND = not determined due to insufficient data

### Massively parallel kinetic measurement of PAI-1 functional half-lives

Leveraging the power of DMS coupled with our phage display system, we developed an approach to accurately extract latency transition half-lives for individual missense PAI-1 mutations from the aggregate half-life data for the entire phPAI-1 mutational library. As described in more detail in *Experimental Procedures*, uPA selection of the library was performed at multiple time points over a 48 h time course (Fig. 1), subjected to HTS, and these data used to determine the half-lives of individual missense mutations. By this approach, we calculated the functional half-lives of 697 individual PAI-1 missense mutations (Fig. 3A, SI Data 2), including 21 of the 35 variants listed in Table 1 (the latter ranging from 2 to 13.1 hours). The mean and median half-lives of the library were determined to be 2.9 and 2.6 h, respectively, and exhibit a strong correlation with functional stability scores (R = 0.6; SI Fig. 1 SI Data 1 and 2).

**Figure 3.**
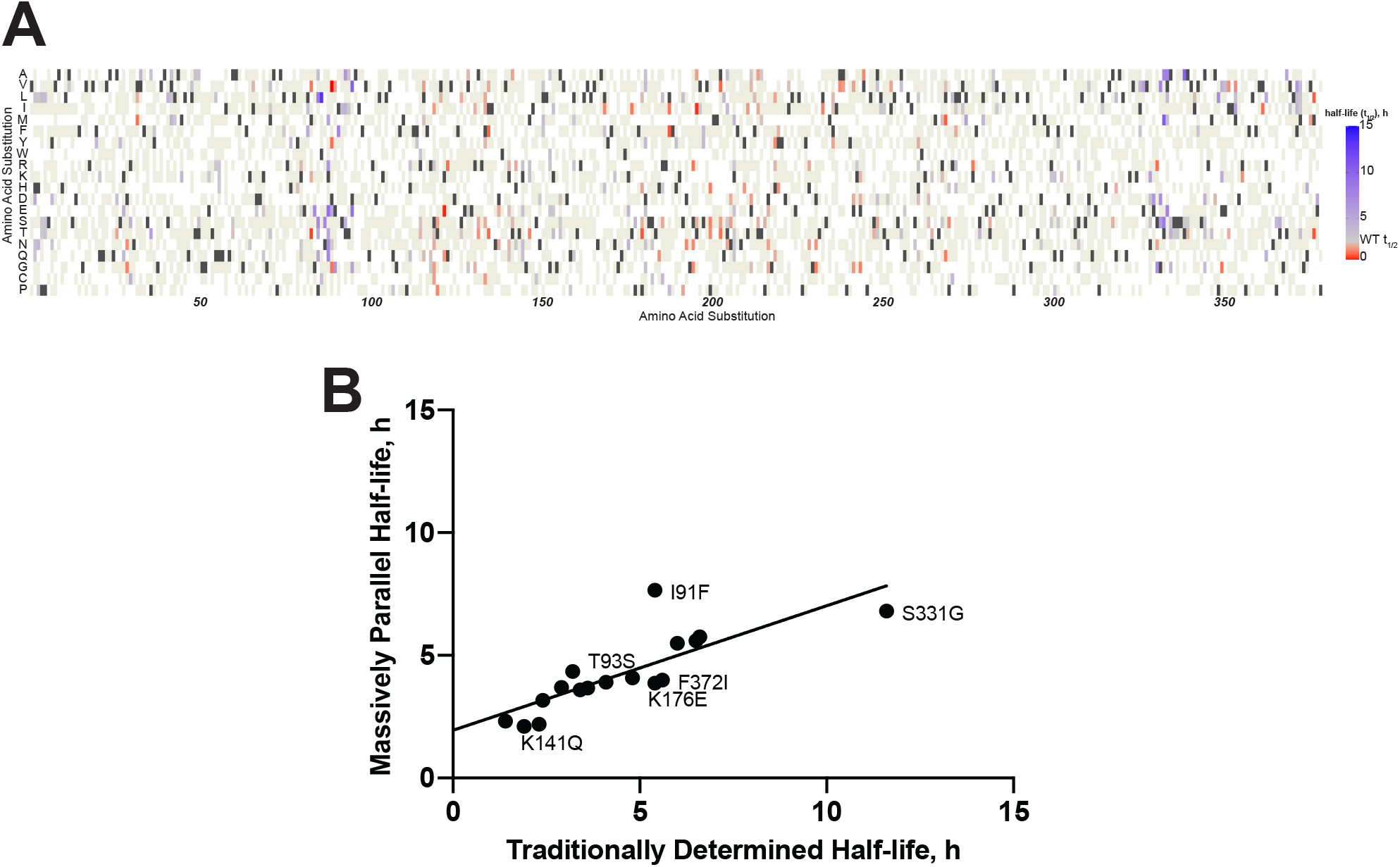
Massively parallel kinetics can accurately determine the half-lives of PAI-1 variants. (*A*) Heatmap indicating average half-lives of phPAI-1 variants. Missense variants with a prolonged half-life (t_1/2_ > 2.1 h) are shown in blue, while those that are decreased are shown in red (t_1/2_ > 2.1 h). WT amino acid residues are indicated by dark grey, while beige indicates missense mutations that were present in the mutational library but for which a half-life different from WT could not be distinguished. (*B*) Half-lives for missense phPAI-1 variants determined using massively parallel kinetics were compared to those described for recombinant PAI-1 both in this study and previously reported (49). A linear regression (slope = 0.5, R = 0.8) indicates a strong positive correlation between our massively parallel method

Additionally, excellent correlation (slope = 0.5, R = 0.8; Fig. 3B) was observed for the half-lives calculated using this massively parallel kinetics compared to those determined for individual rPAI-1 variants in this study (Ile91Phe and Lys176Glu, SI Fig 2) or previously reported (Phe36Thr, Met45Lys, Thr49Ala, Thr93Ser, Lys141Gln, Lys154Thr, Glu212Val, Thr232Ser, Arg271Cys, Gln319Leu, Glu327Val, Ser332Gly, Met354Ile, Phe372Ile, and Phe372Leu) (49).

## DISCUSSION

We report the successful application of DMS to map PAI-1’s mutational landscape with respect to functional stability, identifying a large number of variants that prolong PAI-1’s lifetime in its metastable active or “stressed” state. Though DMS has been increasingly applied to catalog variants for a large number of proteins (50–52), only a few studies to date have attempted to extract kinetic parameters from such large data sets (53–55). The approach described here permits the massively parallel measurement of a kinetic parameter for a large set of amino acid substitutions. A similar approach should be applicable for measuring other parameters including inhibitory rate constants. The results of this and similar studies should also aid in interpreting variants of unknown significance as the clinical use of whole human genome sequencing expands. For example, a PAI-1 variant with a prolonged functional half-life may present with a prothrombotic phenotype, while a PAI-1 variant with a decreased half-life may mimic PAI-1 deficiency (19,20).

In the companion paper to this article, Kihn *et al*. independently applied molecular dynamic simulations and dynamic network analysis to identify residues that facilitate PAI-1’s latency transition, but also those that contribute to an allosteric network of interactions following vitronectin binding, which stabilizes PAI-1 in its active state. Interpreting our DMS data, in the context of the models proposed by Kihn *et al*. suggests that mutations that increase functional stability may do so by disrupting structural features that facilitate the latency transition and/or mimic changes induced by vitronectin binding. For example, the models of Kihn *et al*. suggest that PAI-1’s latency transition is facilitated by a transient salt bridge between Arg^30^ and Asp^193^ (47). Consistent with this hypothesis, while mutations at Asp^193^ are not tolerated (43), we observe that substitutions at Arg^30^ with Cys, His, Leu, or Ser lead to prolonged PAI-1 functional half-lives (4.3, 2.8, 3.4, or 4.5 h, respectively). Given that regions enriched for stabilizing mutations overlap with PAI-1 regions perturbed in response to vitronectin binding, we hypothesize that these mutations may result in a rigidification of the gate-region that mimics that of vitronectin binding (48).

Furthermore, dynamic network analyses identified over 100 trajectories of allosteric pathways that link vitronectin binding to enhanced functional stability. Residues that are degenerate between pathways are either intolerant of mutations (Thr^144^, Tyr^170^, and Met^202^) (43) or the majority of tolerated mutations result in a functionally stable PAI-1 (Leu^169^, Asn^172^, and Asn^329^). These findings suggest that the former residues are not only critical for the identified allosteric networks but also for proper protein folding and function, while the latter stabilizing mutations may replicate or enhance side chain interactions that are induced by vitronectin binding. Our DMS of PAI-1 functional stability identifies several regions in which a large number of amino acid substitutions that increase PAI-1 functional stability are located (Fig. 2, SI Fig. 3). The two regions most enriched in variants that prolong PAI-1’s functional stability are residues 73-98 and 319-347. When mapped to the crystal structure of PAI-1 (PDB: 3Q02 (56)), Fig. 4) residues 73-98 correspond to helix D and β-strand 2A, while residues 319-347 correspond to β-strand 5A and the N-terminus of the RCL (residues 331-350). Other regions enriched in amino acid substitutions that prolong PAI-1’s functional stability include residues 40-49 (helix B), and in the central portion of PAI-1’s primary structure, residues 145-174 (β-strand 2A and the Val^157^-loop).Of note these regions share multiple commonalities to those identified computationally by Kihn *et al*. (48), as well as Jørgensen and colleagues (57).

**Figure 4.**
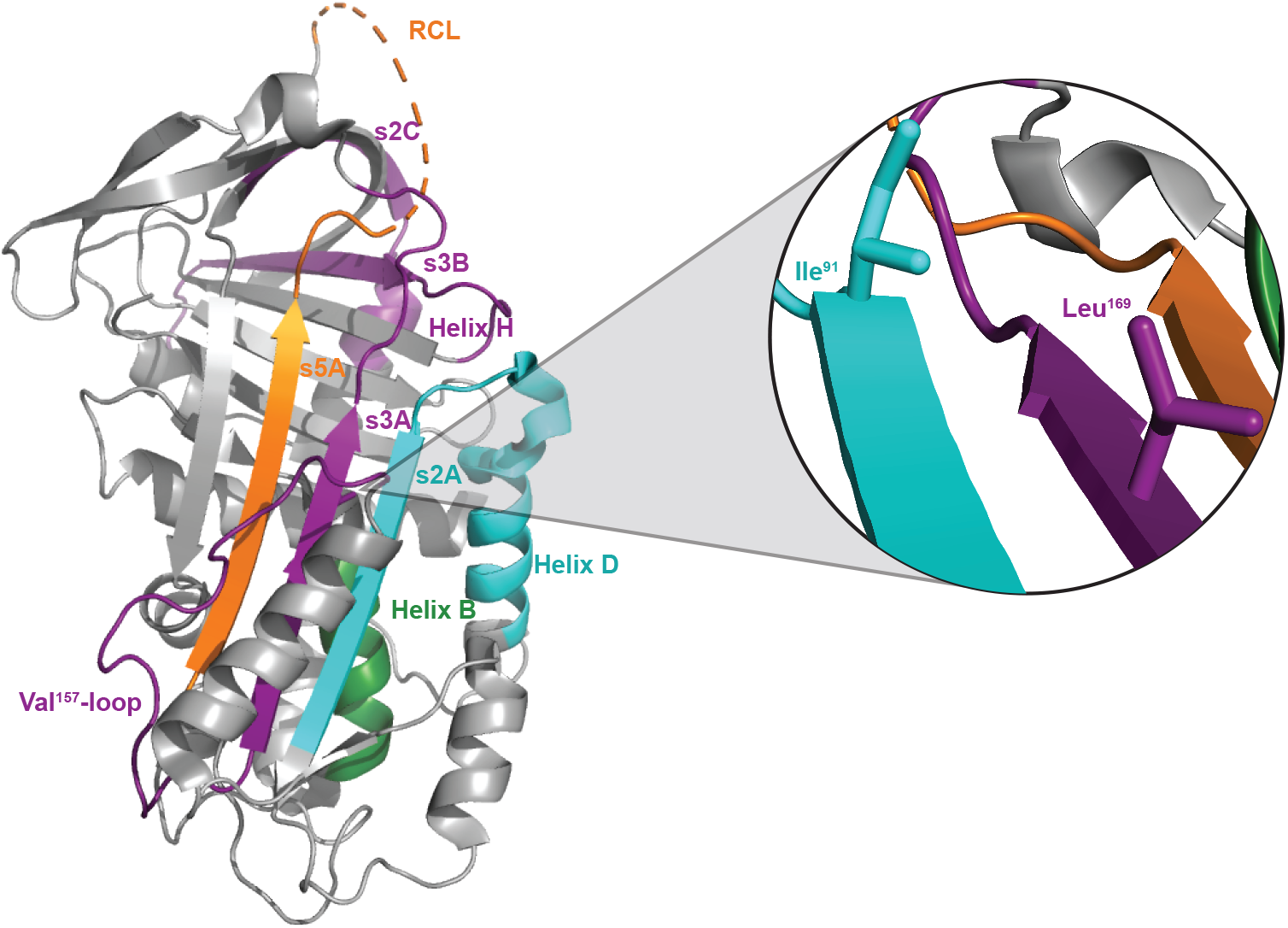
DMS of PAI-1 reveals key regions that regulate its functional stability. The crystal structure of active PAI-1 (PBD: 3Q02 (56)) is shown (*grey*) with regions of increased functional stability relative to WT PAI-1 highlighted with the color coding as indicated in Fig. 3: residues 40-49 map to α-helix B (*green*), residues 73-98 correspond to helix D and β-strand 2A (*cyan*), residues 145-174 encompass the Val^157^-loop and β-strand 3A (*purple*), residues 217-232 correspond to β-strand 2B (*purple*). residues 261-276 encompass helix H and β-strand 2C (*purple*), and residues 319-347 include β-strand 5A as well as the N-terminal portion of the RCL (*orange*). *Inset*, Ile^91^ and Leu^169^ are located in proximity to each other on the interior of β-sheet A (orientation rotated 180° relative to the complete structure).

Helices D and F, along with the Val^157^-loop and the loop connecting helix D and β-strand 2A, are part of one of the most flexible regions of PAI-1, aptly referred to as the “flexible loop region” (27). This flexibility may make it thermodynamically favorable for the F-loop, along with β-strands 1A, 2A, and 3A, to “slide” out of the way as part of PAI-1’s shutter region, permitting full RCL insertion into the central β-sheet A from PAI-1’s pre-latent state (26,58,59). We speculate that the missense mutations in this region identified in our screen most likely decrease flexibility leading to an increased residence time of the RCL in the metastable active and pre-latent conformations.

PAI-1 variants with increased functional stability are also clustered in the “breach region” of PAI-1 between strands 3 and 5 of β-sheet A into which the RCL must insert to undergo either a transition to latency or inhibit a target protease. We hypothesize that mutations in this region may reduce the rate at which the RCL inserts into β-sheet A enough to impede the latency transition but not enough to inhibit the efficient and effective inhibition of uPA (Fig. 4). In a similar vein, the enrichment of mutations in the N-terminal region of the RCL that prolong PAI-1‘s functional stability suggests that latency is driven in part by high-flexibility in this region that allows the RCL to exist in conformations that permit initial insertion into the breach region (Fig. 2).

Other variants of interest that may prolong the functional stability of PAI-1 are clustered in the regions of helices B (residues 40-49) and H (residues 261-276), as well as strand 2 of both β-sheets B (residues 217-232) and C (residues 261-276) as illustrated in Figs. 2 and 4. Flexibility in helix B has previously been linked with an increased probability of a latency transition (27,60). Furthermore, helix H, along with β-strands B and C, comprise the sB/sC binding pocket of PAI-1 in which ligands can allosterically inhibit vitronectin binding to PAI-1 (61). We postulate that mutations in this region that extend the functional half-life of PAI-1 may mimic the allosteric changes in PAI-1 structure induced by vitronectin binding, potentially restricting the RCL from transitioning to a latent state, while not rendering PAI-1 so rigid that it loses its inhibitory capacity.

Of the 35 most enriched mutations listed in Table 1, two or more different amino acid substitutions were observed at seven distinct sites (Ile^91^ (n = 3), Leu^169^ (n = 2), Asp^222^ (n = 4), Tyr^241^ (n = 2), Gly^332^ (n = 3), Thr^333^ (n =2) and Phe^372^ (n = 4)). Among these sites, Gly^332^ and Thr^333^, along with Ser^331^, are located at the N-terminal hinge of the RCL. Mutation of Gly^332^ and Thr^333^ to other small amino acids (Ser, Asp, and Ala for Gly^332^, as well as Ala and Ser for Thr^333^) may not affect RCL insertion into β-sheet A while moderately affecting the accessible Ramachandran space, thereby increasing the time scale of RCL insertion—consistent with limiting the ability of PAI-1 to enter a pre-latent state (26).

Residues Asp^222^ and Tyr^241^ are located on the periphery of β-sheet B. Sui and Wiman have previously reported that mutations in this region are linked to prolonged functional stability of PAI-1 (62). Although their mutational analysis suggests that substitution of a basic residue (Lys) at position 222 is responsible for the change in functional stability, our more comprehensive mutational mapping demonstrates that simple loss of an acidic amino acid at this position by mutation to a neutral side chain (His, Tyr, Val, Asn) may be more critical. Sui and Wiman also reported that Tyr241Ser resulted in a modest increase in PAI-1 functional stability, consistent with our data for multiple substitutions at this site (Table 1; SI Data 1 and 2).

Three other amino acid positions Ile^91^, Leu^169^, and Phe^372^ with multiple enriched mutations are clustered on strands 2 and 3 of β-sheet A (Table 1). Previously, the Ile91Leu mutation was reported as the single amino acid substitution that most significantly prolongs the half-life of PAI-1 (44). The identification of two additional amino acid substitutions at this position that increase the functional stability of PAI-1 (Ile91Phe and Ile91Met) further highlights the importance of this position and suggests that the presence of bulkier side chains stabilizes the shutter region in a closed position. Likewise, two of the four mutations at Phe^372^ (Phe372Ile and Phe372Leu) had been previously identified to stabilize PAI-1 in its metastable state. The additional two variants identified here (Phe372Val and Phe372Cys) further suggest that substitution of Phe with a smaller hydrophobic residue promotes PAI-1’s functional stability. Though the importance of Leu^169^ in affecting PAI-1’s functional stability has not been reported previously, our data identify both Leu169His and Leu169Ser as promoting enhanced stability in PAI-1’s active stressed state—perhaps due to increased potential for hydrogen bond formation or hydrophobic interactions, as Ile^91^ and His^169^ make contact with each other on the interior of β-sheet A (Fig. 4, *inset*). As noted above, Kihn *et al* (48) also identify Leu^169^ as a key residue in predicted allosteric networks that drive PAI-1 functional stability following vitronectin binding.

(48)

The application of high-throughput kinetics to accurately map PAI-1 function highlights a novel way in which to couple traditional phage display with DMS technology. The phPAI-1 mutational library used in these studies contains multiple amino acid substitutions per molecule, and thus epistatic interactions between mutations in cis with each other on different sequencing amplicons (SI Fig. 4) may interfere with the determination of quantitative half-lives in these studies. However, the excellent agreement between our massively parallel calculations and the values measured for individual variants suggests that this factor does not significantly confound these data (Fig. 3B). Similar to the conclusions of Kretz *et al*. in their massively parallel kinetic studies of the rates for ADAMTS13 proteolysis of a phage displayed peptidyl library (55), we posit that any epistatic effects average out over multiple clones, generating a neutral background.

We fully anticipate that a similar approach could be applied to other high-throughput kinetic assays (63,64), including those that occur on faster time scales (e.g. inhibition rate constants), as catalytic efficiency has been previously measured using a related method (64). Of note, in the present study the experimental design was established to identify those PAI-1 variants with high functional stability. Even then, we identified an inherent issue with calculating massively parallel half-lives in this manner—for the largest functional enrichment scores (Table 1), the massively parallel half-lives are underrepresented and are the least accurately predicted by their associated functional stability score (SI Fig. 1). As has been demonstrated by others (54), altering the reaction conditions to compensate for the wide range of possible kinetic parameters should ameliorate this experimental artifact.

In conclusion, we have applied DMS to map the mutational landscape of PAI-1 with respect to its latency transition. Overall, we confirm and identify structural features that affect the functional stability of PAI-1 that complement not only prior experimental work but also molecular dynamics simulations of PAI-1 behavior (65), highlighting the utility of high-throughput unbiased screens for protein variant function. We were also able to accurately and quantitatively determine the half-lives of PAI-1 variants across its mutational landscape, allowing biochemically meaningful value to be incorporated into our high-throughput screen. DMS combined with enzymatic and kinetic measurements has the potential to couple biochemistry, structural biology, and genomics in which the unbiased screens provide a unique understanding of the complex systems at play in both protein evolution and protein architecture.

## EXPERIMENTAL PROCEDURES

### Generation of phage display library

WT phPAI-1 and the phPAI-1 library were generated, characterized, and expressed as previously described (43). Briefly, WT human SERPINE1 cDNA with an N-terminal *myc*-tag and Gly-Gly-Gly-Ser linker was cloned between the AscI and NotI restriction sites of pAY-FE (Genbank #MW464120) (34,66). The phPAI-1 library was generated by error prone PCR using the GeneMorph II Random Mutagenesis Kit (Agilent Technologies, Santa Clara, CA) with primers that maintain the AscI and NotI restriction sites (SI Table 1). Following cloning of the phPAI-1 library into the pAY-FE vector, the library was transformed into electrocompetent XL-1 Blue MRF’ *E. coli*. The resulting library has a depth of 8.04×10^6^ independent clones with an average of 4.2 ± 1.8 amino acid substitutions per PAI-1 fusion protein (43).

As previously reported (43,66), phage were produced by growing *E. coli* harboring pAY-FE PAI- −1 in LB broth supplemented with 2% glucose and ampicillin (100 μg/mL) at 37 °C to mid-log phase and infecting with M13KO7 helper phage for 1 h. Bacteria were subsequently transferred to 2xYT media supplemented with ampicillin (100 μg/mL), kanamycin (30 μg/mL), and IPTG (0.4 mM) to induce phPAI-1 expression for 2 h at 37 °C. All subsequent steps were carried out at 4 °C to minimize the PAI-1 latency transition. Phage were precipitated with polyethylene glycol-8000 (2.5% w/v) and NaCl (0.5 M) for 2-16 h followed by centrifugation (20,000xg for 20 min at 4 °C) (67). The precipitated phage pellet was resuspended in 50 mM Tris containing 150 mM NaCl (pH 7.4; TBS).

### phPAI-1 latency screen

phPAI-1 (100 uL at ~10^12^ phage/mL, ~0.17 nM final concentration) were incubated in TBS containing 5% BSA (900 μL) at 37°C and aliquots assayed over a 48 h time course (0, 1, 2, 4, 8, 12, 24, 48 h). At each time point, uPA (1.7 nM) was incubated with the phage for 30 min at 37°C. With an estimated valency of less than one PAI-1 fusion protein per phage (43), this is equivalent to at least a 10-fold excess of uPA. Residual uPA activity was inhibited by the addition of 100 uL of a 10X stock of cOmplete EDTA-free protease inhibitor cocktail (Millipore Sigma) and incubated for 10 min at 37 °C. phPAI-1:uPA complexes were further immunoprecipitated, washed, eluted, and quantified as described previously (25). In parallel experiments, phPAI-1 was immunoprecipitated via its N-terminal myc-tag over the same time course (Figs. 1).

### High-throughput sequencing (HTS)

Twelve overlapping amplicons (150 bp) were PCR amplified from pAY-FE PAI-1 (SI Table 1 and SI Fig. 4) and prepared for HTS as previously described (43). HTS data were analyzed using the DESeq2 software package (68). Statistical thresholds (padj < 0.05 and base mean score > 10) were used to classify a mutation as significant for its effect on functional stability, where padj is the minimum false discovery rate when calling a mutation significant and the base mean score is the mean of the normalized counts across all samples. Missense mutations in PAI-1 that were nonfunctional were removed from subsequent analyses (43), and a functional stability score was calculated for each mutation by subtracting the log2-fold change (uPA-selected over unselected) for WT PAI-1 from that of each amino acid substitution at each position.

### Recombinant PAI-1 (rPAI-1)

Human *SERPINE1* cDNA was subcloned from pAY-FE-PAI-1 using ligation independent cloning into the pET Flag TEV LIC cloning vector (2L-T, a gift from Scott Gradia—Addgene plasmid #29175; http://n2t.net/addgene:29715; RRID:Addgene_29715) using primers pET PAI-1 LIC For and pET PAI-1 LIC Rev (SI Table 1) and transformed into BL21(DE3) chemically competent *E. coli* (Invitrogen) (69). Point mutations (I91F and K176E) were introduced using the QuickChange II XL Site-Directed Mutagenesis Kit (SI Table 1; Agilent Technologies). The full-length sequence of the PAI-1 constructs was confirmed by Sanger sequencing (SI Table 1). For recombinant PAI-1 (rPAI-1) expression and purification, *E. coli* containing pET-PAI-1 constructs were grown at 37°C in LB media (100 mL; Invitrogen) supplemented with ampicillin (100 μg/mL) to mid-log phase (OD_600_ ~0.40-0.45) prior to induction with IPTG (1 mM) and continued growth for 2 h at 37 °C. Bacteria were pelleted by centrifugation (3500xg for 20 min at 4°C) and washed twice with NaCl (0.85M). The bacterial pellet was resuspended in sodium phosphate (50 mM) pH 7.0 containing NaCl (100 mM) and EDTA (4 mM) and subsequently incubated with T4 lysozyme (1 mg/mL; Millipore Sigma) for 30 min at ambient temperature. Following three freeze-thaw cycles (1 min in liquid nitrogen, 5 min at 37°C), the bacterial genomic DNA was sheared by passing 5-10 times through a 20-gauge needle, and crude cell extracts isolated following centrifugation (16,000xg for 20 min at 4°C). rPAI-1 was further purified by binding to anti-FLAG agarose beads and elution with 3X FLAG peptide (Millipore Sigma). rPAI-1 was quantified spectrophotometrically (ε_280nm_ = 43,000 M^−1^ cm^−1^). To measure its functional half-life, rPAI-1(20 nM) was incubated at 37°C over a 24 h time course and assayed for activity. At selected time points, the rPAI-1 was diluted between 2- and 40-fold in HEPES (40 mM) pH 7.4 containing 100 mM NaCl and 0.005% Tween-20 and its inhibitory activity against uPA (2.5 nM) was measured following incubation for 30 min at room temperature. Residual uPA activity was assayed by the rate of proteolysis of the fluorescent substrate Z-Gly-Gly-Arg-AMC (Bachem) over a 10 min time course (λ_ex_ = 370 nM, λ_em_ = 440 nM) (SI Fig. 2).

### Transition to latency half-life calculation

The functional half-lives of both phPAI-1 and r PAI-1 were determined by plotting either the fraction of phPAI-1 recovered or relative activity of rPAI-1 that remains active after fitting with a one-phase exponential decay function (Eq.1)

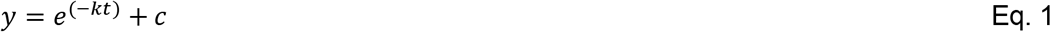

where y is the PAI-1 fractional activity, k is the rate constant for the first-order transition of PAI-1 from its active to latent form, t is time, and c is the asymptotic value. Half-life (t_1/2_) was further determined as described in Eq. 2.

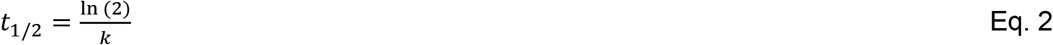

### Massively parallel half-life kinetics

To determine the half-lives of individual PAI-1 missense variants within the library, the fractional activity of the library at each time point over a 48 h time course (Fig. 1) was multiplied by the fraction of counts representing a given missense mutation that were present at each time point. Following normalization of the 0 h time point to 1, the data were fit to an exponential decay function (Eq. 1) and the apparent half-life extracted (Eq. 2). In order to account for variations in the mutational background arising from error prone PCR, the variants at a given position were normalized by setting the WT amino acid substitution to half-life of 2.1h.

## DATA AVAILABILITY

All data is contained within the manuscript and supplementary files. Bioinformatics scripts are available at https://github.com/hayneslm/PAI-1_functional_stability. Raw sequencing data will be made available upon request.

## SUPPORTING INFORMATION

This article contains supporting information (4 Figures, 1 Table, and 2 Data Files).

## FUNDING AND ADDITIONAL INFORMATION

This work was supported by the National Institutes of Health grants R35-HL135793T (DG),R01-HL055374 (DAL), R01-AG074552 (DAL) T32-GM007863 (ZMH), and T32-HL125242 (ZMH). LMH was supported by a National Hemophilia Foundation Judith Graham Pool Postdoctoral Research Fellowship. AY is supported by an American Society of Hematology Faculty Scholar Award, National Hemophilia Foundation Innovative Investigator Research Award, National Institutes of Health grant R01-HL154688, and the Mary R. Gibson Foundation. CAK is supported by an Early Career Award from Hamilton Health Sciences, the Canadian Institutes of Health Research (PJT-168987), and the Natural Sciences and Engineering Research Council of Canada (DG-5007801). DAL has additional support from the American Heart Association (19TPA34880040). DG is a Howard Hughes Medical Institute investigator. The content is solely the responsibility of the authors and does not necessarily represent the official views of the National Institutes of Health.

## ACKNOWLEDGEMENTS

We thank Kyle Kihn, Elisa Marchiori, Giovanni Spagnolli, Alberto Boldrini, Luca Terruzzi, Daniel Lawrence, Anne Gershenson, Pietro Facciolo, and Patrick Wintrode for access to their computational models prior to publication.

## ABBREVIATIONS

DMS: deep mutational scanning
HTS: high-throughput sequencing
PAI-1: plasminogen activator inhibitor-1
phPAI-1: phage displayed PAI-1
RCL: reactive center loop
SERPIN: serine protease inhibitor
t_1/2_: half-life
tPA: tissue-type plasminogen activator
uPA: urokinaselike plasminogen activator
WT: wild type

