## Supplementary material for "Deep mutational scanning and massively parallel kinetics of plasminogen activator inhibitor-1 functional stability": SI Fig

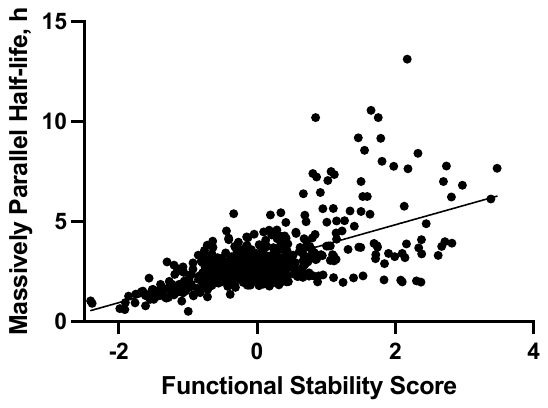

**SI Figure 1. Functional stability scores positively correlate with massively parallel half-lives.** A positive correlation (R = 0.6) was observed between the functional stability scores and the massively parallel half-lives.

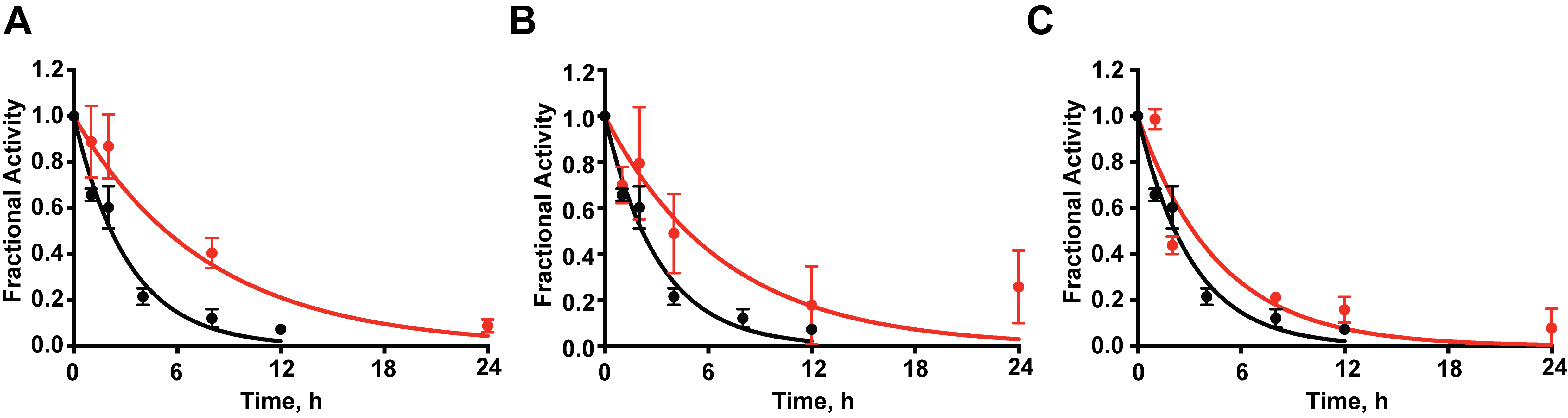

**SI Figure 2. Rates of latency transition for recombinant PAI-1 variants *(A)* I91F and *(B)* K176E.** Activity for WT rPAI-1 is indicated in black and single amino acid substituted variants are shown in red (n = 3). The half-life of I91L rPAI-1 was determined to be 5.4 h, while for K176E it was 4.8 h. WT rPAI-1 had a half-life of 2.2 h.

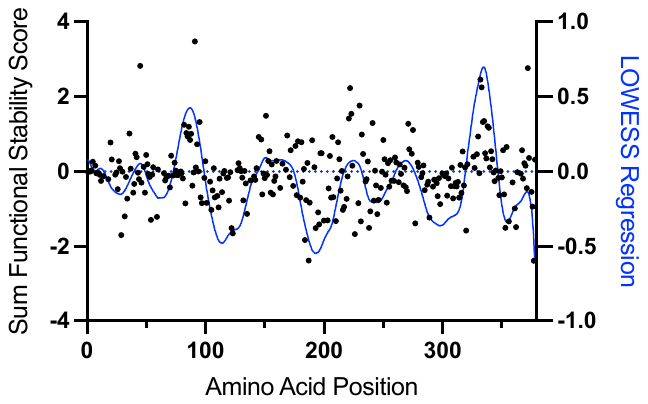

**SI Figure 3. Identification of PAI-1 regions enriched in variants that stabilize the active conformation.** PAI-1 amino acid positions are plotted on the x-axis with black circles representing the sum of functional stability scores divided by the number of variants scored at each position The data were fit to a LOWESS regression (*right* axis, blue) with a 20 point smoothing window determined with the GraphPad Prism software package (v. 9.0.2). Six regions of increased functional stability were identified with this method (LOWESS regression > 0): residues 40-49, 73-98, 145-174, 217-232, 261-276, and 319-347.

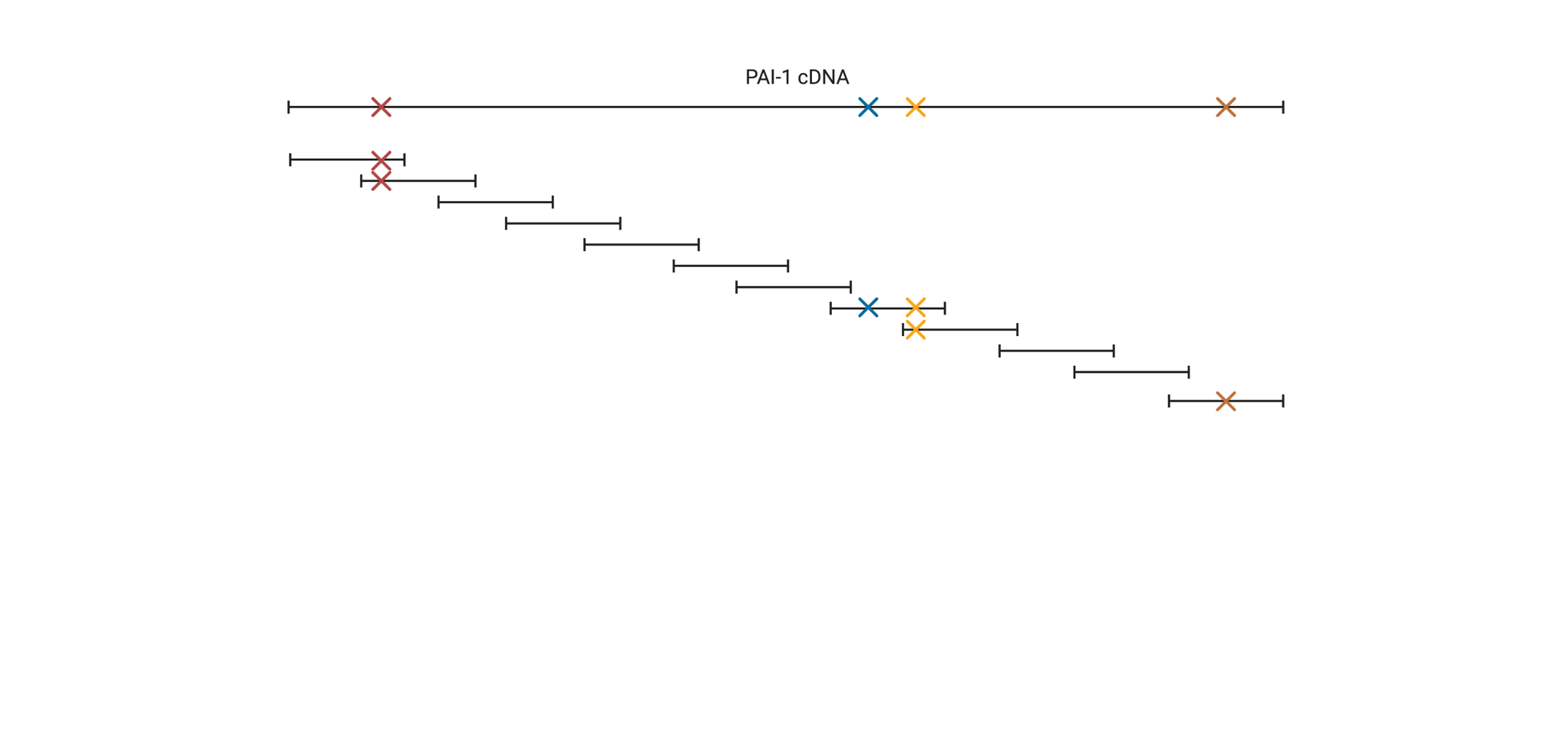

**SI Figure 4. A hypothetical mutated PAI-1 cDNA is shown with 4 mutations.** Mutated PAI-1 cDNAs are sequenced using 12 overlapping 150 bp amplicons. Most amplicons will be WT with no mutations. Most mutations will only appear in 1 amplicon (blue and brown) and phase will be lost with mutations in *cis* in the original cDNA but occurring in different amplicons. (Created with BioRender.)
